# A Systems Change Framework for Evaluating Academic Equity and Inclusion in an Ecology & Evolution Graduate Program

**DOI:** 10.1101/848101

**Authors:** Kelly J Wallace, Julia M York

**Author notes:** both authors contributed equally. Authors for correspondence: Kelly J. Wallace & Julia M York, University of Texas at Austin, Department of Integrative Biology, 1 University Station C0990, Austin, Texas 78712.

## Abstract

While academia is moving forward in terms of diversifying recruitment of undergraduate and graduate students, diverse representation is still not found across the academic hierarchy. At the graduate level, new discussions are emerging around efforts to improve the experiences of women and underrepresented minorities through inclusive graduate programming. Inclusive graduate programs are that which actively center and prioritize support for diverse experiences, identities, career goals, and perspectives, from recruitment through graduation. Establishing regular and rigorous evaluation of equity and inclusion efforts and needs is a critical component of this work. This is recognized by funding agencies that increasingly require reporting on inclusion efforts; here we suggest use of a systems change framework for these evaluations.

A systems change approach emphasizes three levels: explicit change (e.g. policies), semi-explicit change (e.g. power dynamics), and implicit change (e.g. biases). We use the Ecology, Evolution, and Behavior (EEB) PhD Program at the University of Texas at Austin in an exercise to (1) identify areas of concern regarding inclusive programming voiced by graduate students, (2) categorize efforts to address these concerns, and (3) integrating and evaluating which areas of the systems change framework show the greatest progress or potential for progress. We argue this framework is particularly useful for academic systems as they are complex, composed of variable individuals, and must address diverse stakeholder needs.

## General Background

Since 2008, women have earned 60% of baccalaureate degrees and the majority of doctoral degrees in biology in the U.S., and in 2016, 42% of baccalaureate degrees and 32% of doctorates in biology were earned by underrepresented minorities (NSF, 2019). Thus, the elementary demographics of biology doctorate earners is roughly representative of the U.S. population as a whole, which was 51% women and 36% non-white in 2017 (although much intersectional work remains beyond these most crude categorizations; Patridge et al., 2014; Van Cooten, 2014; Li and Koedel, 2017; Reardon, 2017; Brown and Leigh, 2018; NSF, 2019; Barnes et al., 2020). This representation, however, is dramatically reduced at the faculty level: only 35% of biology faculty are women and 25% are people of color (among full professors 15% are people of color; NSF, 2019). If faculty demographics were representative of the available PhD applicant pool, we are living in 1987 (the most recent year when women accounted for 35% of biology doctorate earners; NSF, 2017). So why is biology academia more than 30 years behind?

We suggest that experiences in graduate school are a determining factor of the leaky pipeline. During graduate school, most PhD students first experience and internalize the academic lifestyle, and the majority choose another career path (Joseph, 2012; Mack et al., 2013). While a successful PhD program prepares students for the myriad careers doctorate holders in biology eventually pursue (Turk-Bicakci et al., 2014), those leaving academia are disproportionately women (Martinez et al., 2007; Shaw and Stanton, 2012; Glass et al., 2013) and underrepresented minorities (Allen-Ramdial & Campbell, 2014; Li and Koedel, 2017; **Figure 1A)**; this must change. There is not one single reason why women and underrepresented minorities leave academia. Rather, it is a comprehensive and nuanced set of experiences wherein marginalized students use the acumen and perception that gained them acceptance into their doctoral program to learn the many ways in which the academic system is not built for them (Ong et al., 2011; Puritty et al., 2017; Slay et al., 2019; Makarem and Wang, 2020). This is a product not only of their own experiences, but in also the keenly observed experiences of co-workers and representative faculty (Settles et al., 2006; Case and Richley, 2013; Hirshfield, 2014; Patridge et al., 2014; San Miguel and Kim, 2015; Yoder and Mattheis, 2015).

**Figure 1.**
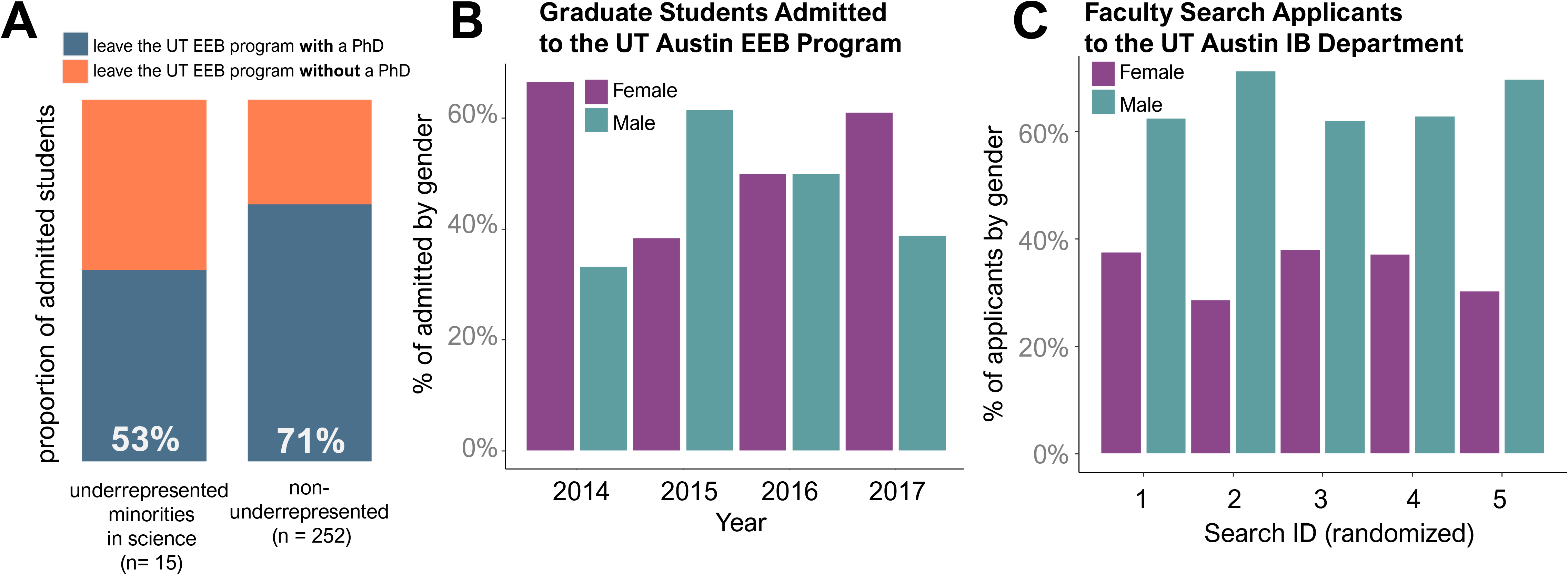
Demographic representation statistics for the Ecology, Evolution, and Behavior Graduate Program and the Department of Integrative Biology at the University of Texas at Austin. (A) A comparison of PhD graduation rates of under-represented (Black, Hispanic, and Native America) and non-underrepresented admitted students (white and Asian) in the EEB Program. (B) Gender demographics of graduate students admitted into the PhD Program from 2014-2017 and (C) gender demographics of applicants to the IB Department’s five most recent faculty searches (randomized for anonymity).

Many graduate programs in biology have implemented spaces, techniques, conversations, and policies to improve graduate student well-being, particularly for under-represented students (Bekki et al., 2013; Porter et al., 2018; Williams and Korn, 2018; Gold et al., 2020). Even as these individual efforts are successful, however, the inequities they attempt to counter persist. Why? We posit that reliance on unimodal or sporadic diversity efforts is insufficient for fundamental change. Instead, programmatic diversity efforts must be orchestrated to sustain systemic change. We propose using a *systems change framework* to critically evaluate, develop, and coordinate equity and inclusion efforts across system modes (Coffman, 2007). A systems change framework (described in more detail below) is not a new concept, but here we argue that it is a particularly useful framework for biology graduate programs to critically evaluate the cross-hierarchical, systemic challenges and reform efforts that graduate programs face and implement, respectively, particularly as it relates to mentorship, diversity, and inclusion.

Here we detail an exercise for graduate programs to evaluate and strategize their inclusivity efforts. In undertaking the exercise for our graduate program as an example, we demonstrate the value of a systems change framework in identifying the areas of progress and areas of need in a graduate program. Using a systems change framework, we categorize the most common gaps in support for graduate students, allowing us to “see the water” of the system, and then we categorize recent programmatic efforts to address those concerns (Kania et al., 2018). We use this paired framework to illuminate areas of progress and areas in need of increased focus for future program development.

As biology PhD candidates ourselves, we lack direct power to enact changes that would reform the system on an institutional level. Availability of mental health resources, handling of harassment and misconduct cases, selection of administration, and family leave policies are all in dire need of systemic reform by those with administrative power (Anders, 2004; Handelsman et al., 2005; Case and Richley, 2013; Su et al., 2015). In addition to advocating for these institutional reforms, we believe we can counteract negative graduate student experiences by focusing on departmental culture and climate reform, utilizing thoughtful and inclusive data collection on the quality of the graduate student experience followed by implementation of actions based on those data (Institute of Medicine, 2007; Tao and Gloria, 2019, Slay et al., 2019). We include more information on our individual inclusivity efforts in the supplemental materials for those interested, but our goal here is not to summarize or elaborate on those efforts, it is rather to engage in an exercise where we critically evaluate them in the context of the systemic issues they were implemented to address.

### What is a Systems Change Framework?

The theory of *systems change* is designed to reform the underlying conditions in a system as they relate to social change, diversity, and inclusion, and was originally conceived in activist pedagogy (Coffman, 2007). Its early applications centered around access to resources related to physical and mental health in early childhood development, and recently has become more frequently utilized in corporate management areas and social organizations (Kania et al., 2018; Seelos and Mair, 2018). The systems change framework is a construct intended to organize and evaluate the needs and corresponding efforts of a community, especially when that community is composed of diverse stakeholders. It allows the community system to become the focus of inquiry, rather than individual victims or perpetrators (Foster-Fishman and Watson, 2018). This makes it particularly useful for academic systems where stable conditions result from variable and dynamic individuals and individual actions (Jenal and Cunningham, 2020).

The systems change framework itself is a descriptive set of interconnected spheres or categories of influence of a “system” - for example, a program, department, school, business, organization, or initiative. Systems operate on many organizational levels (e.g. individual, community, state), often have a variety of funding sources, and must “tackle difficult deep-rooted problems such as gaps in services and outcomes based on race, income, culture, and language” (Coffman, 2007). The framework allows these complexities to be dissected and evaluated without losing sight of the system as a whole (Foster-Fishman and Watson, 2018). Systems frameworks do this by emphasizing understanding the system and tailoring interventions, rather than focusing on success or failure of individual efforts (Seelos and Mair, 2018). Given this, we believe the framework provides a clear format to help graduate programs evaluate and tackle such deep-rooted and complex problems as persist in academia.

The literature on systems change varies in nomenclature and the number of categories, here we choose to utilize the framework described in Kania, Kramer, and Senge (2018) “The Water of Systems Change.” Kania et al. highlight six “conditions” or areas of systems change that fall into three categories: explicit, semi-explicit, and implicit. We adjust the definitions to the six conditions used by Kania et al in terms of specificity to graduate programs **(Figure 2)**.

**Figure 2.**
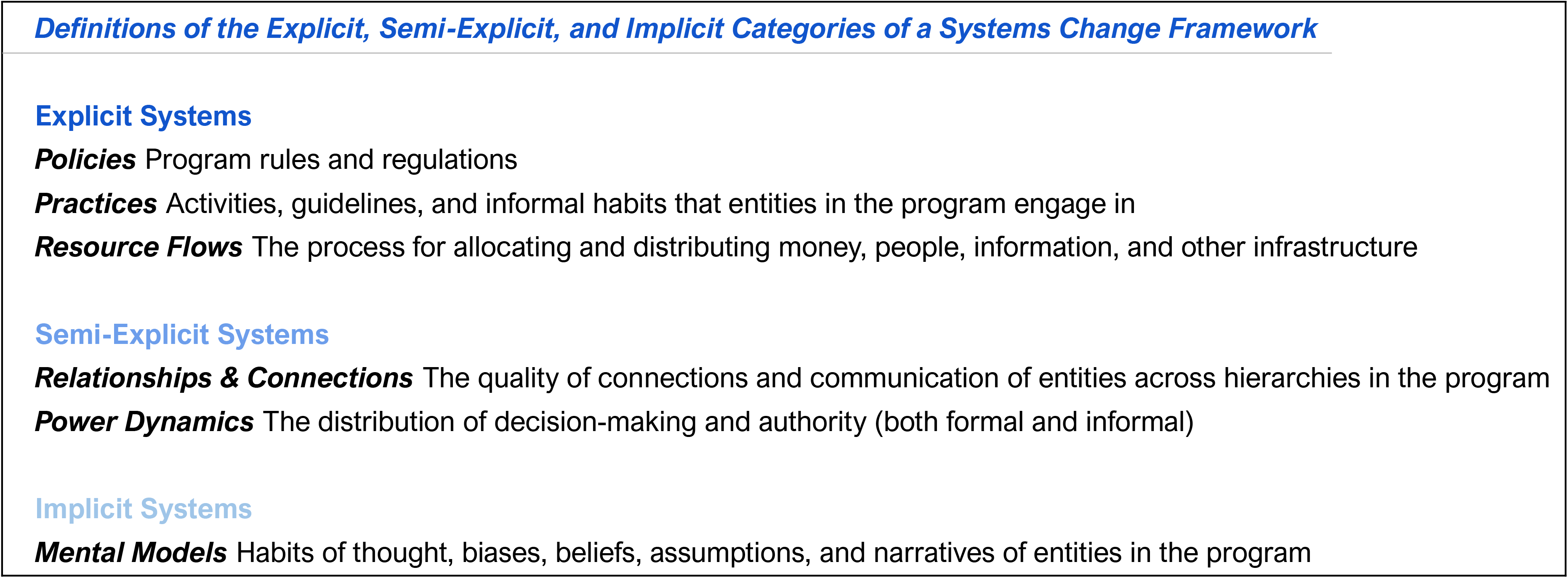
Definitions of the explicit, semi-explicit, and implicit categories of a systems change framework, adapted for biology academic graduate programs

### Exercise General Description

We designed an exercise that consists of three primary parts:

Step 1: *Data collection/system assessment: Regularly assess the most common concerns or needs expressed by all stakeholders in the program*
Step 2: *Identify currently existing or proposed efforts developed to address those concerns/needs. Organize those efforts into the categories of the systems change framework*
Step 3: *Evaluate the areas of overlap and limitations from Steps 1 and 2. Identify which categories of the systems change framework show the greatest progress in tackling concerns, and conversely which categories show the most urgent need for additional attention*

In the following sections we apply this to our graduate program: the Ecology, Evolution, and Behavior (EEB) Graduate Program at the University of Texas at Austin, as an example. We believe this program is an effective example because we are addressing challenges present in other programs and our demographics are roughly similar to national averages (Princeton Graduate Women in STEM Leadership Council, 2018; Slay et al., 2019; NSF, 2019). Current graduate students are 52% female while supervising faculty are 34% female (22% of the senior faculty). This disparity in representation is reflected across several pools of data: for example, comparing the gender of the admitted graduate students with that of the faculty applicant pool **(Figure 1B, 1C)**. Data regarding racial and ethnic composition is more difficult to obtain due to small sample sizes, as well as being more difficult to disseminate while protecting anonymity, yet show that admitted students from minority groups underrepresented in science (Black, Hispanic, and Native American) show lower PhD graduation rates **(Figure 1A)**. We recognize that the information we present here still does not fully encompass all the unique and intersectional challenges that many other underrepresented groups face (e.g. sexuality, disability status, socioeconomic status) and see this as a strong avenue for additional work.

### Step 1: Identifying and categorizing areas of concern through data collection

We collected a list of common concerns expressed by graduate students in the program. This list was primarily comprised of responses to a comprehensive climate survey developed in 2018 and administered for the department by one of the authors (supplemental material 2) and was supplemented by the first-hand experiences of the authors and conversations regarding experiences of other students. We acknowledge this list is not exhaustive and may be limited based on our own mental models and personal relationships, however we believe the practice of compiling this list builds capacity to understand and “see” the program as a system. In describing the items on the list, we took great care to keep information as anonymous as possible. We then sorted these concerns into the categories of the systems change framework (explicit, semi-explicit, implicit) as well as specific-sub-categories **(Figure 3)**.

**Figure 3.**
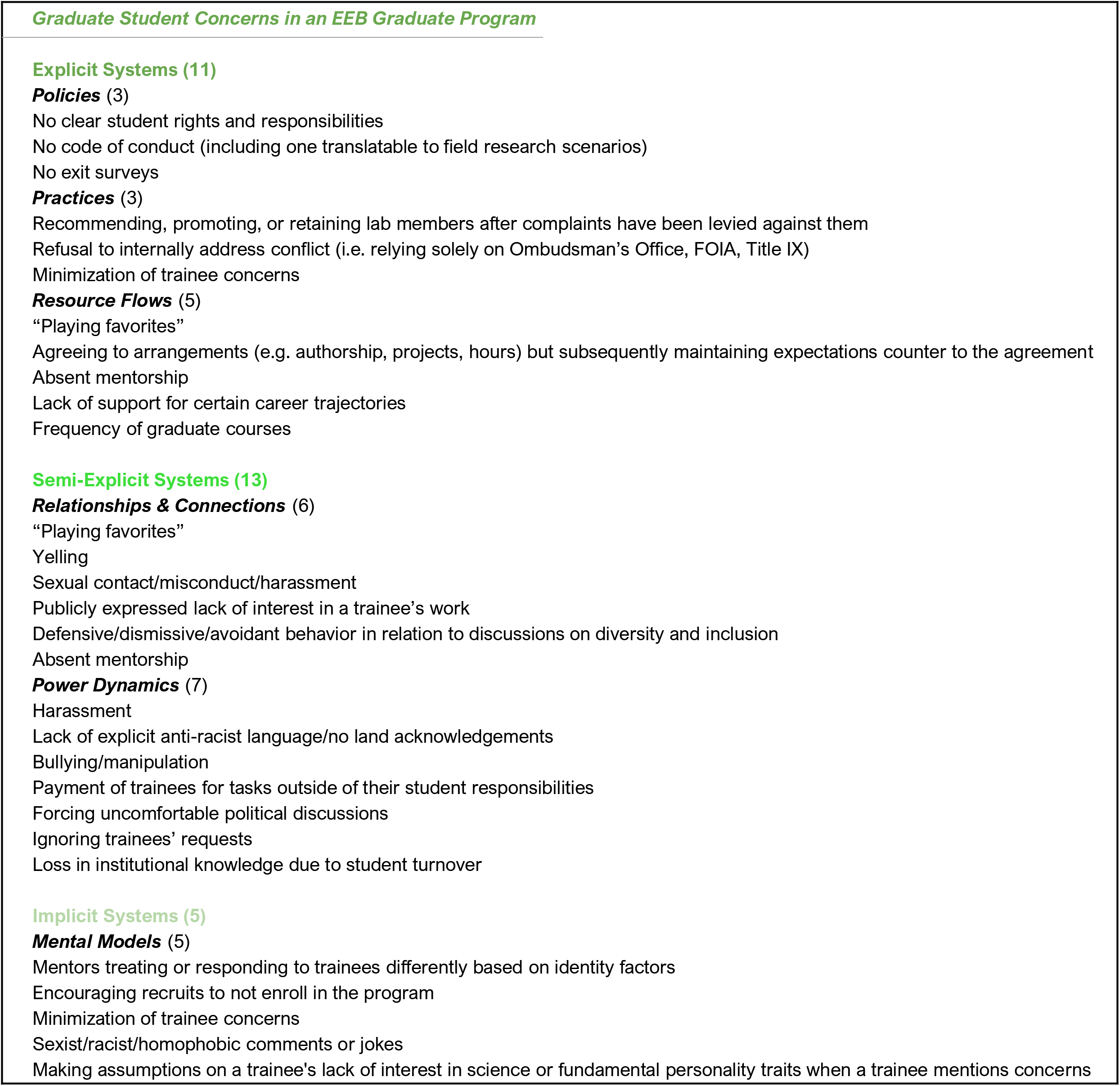
Graduate Student Concerns in an EEB Graduate Program

### Step 2. Identifying Efforts

Students in the UT Austin EEB Graduate Program have spent significant time and energy developing and implementing reforms to address many of the concerns described in Step 1. We listed these efforts, categorizing them into the systems change framework **(Figure 4)**. Here we do not elaborate further on the details and value of each individual effort, though more information regarding these efforts can be found in supplemental materials or by contacting the authors. For the purposes of this exercise we focus on which *categories* of the systems change framework each effort falls under to identify broader patterns of need and progress.

**Figure 4.**
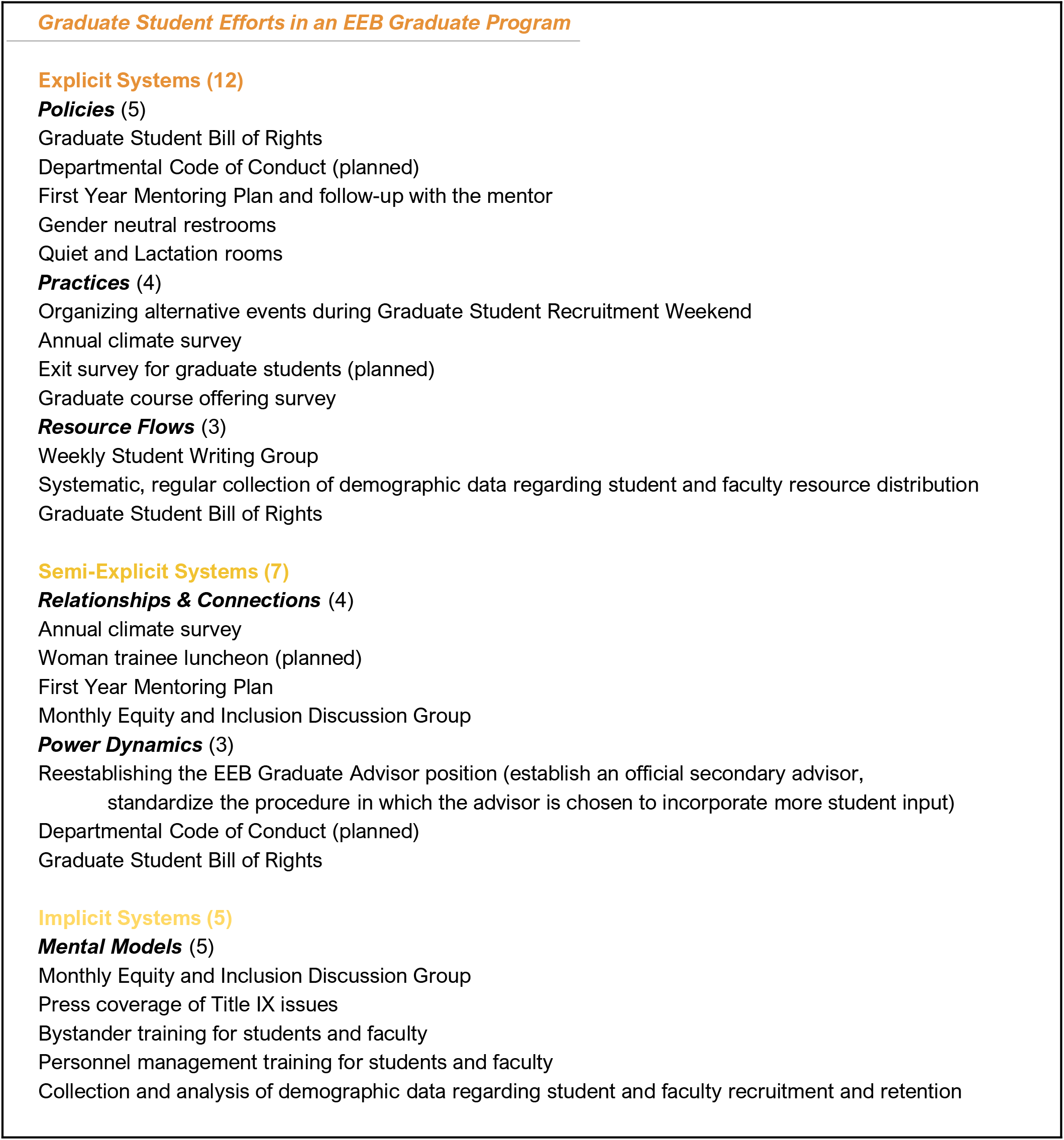
Graduate Student Efforts in an EEB Graduate Program

### Step 3. Evaluating Areas of Greatest Progress and Greatest Need

We evaluated the areas of greatest progress and need by asking four questions: (1) which systems change category housed the largest number of concerns, (2) which category housed the largest number of efforts, (3) in which category did the concerns most “outweigh” the efforts, and (4) in which category did the efforts most “outweigh” the concerns? We find the largest number of concerns were within semi-explicit systems, while the majority of efforts were focused on explicit systems. Therefore, semi-explicit concerns most outweighed the existing efforts in that category, whereas efforts to tackle explicit systems were more frequent than concerns expressed about explicit systems. This assessment guided our further analysis of our program as a system.

## Discussion

We have described a three-step exercise that graduate programs can engage in to evaluate their equity and inclusion efforts and identify areas of greatest progress and greatest need. When undergoing this exercise for the home program of the authors (the University of Texas at Austin EEB Graduate Program), we find that student concerns primarily fall into the semi-explicit category of the systems change framework (Relationships & Connections / Power Dynamics).

In this EEB program, graduate students in general did not express strong concern regarding explicit policies--we evaluate this to mean that the program is successfully implementing policies that support a diverse and inclusive climate. Collaboration, infrastructure, variety of expertise, grant and fellowship success, and publication quality were not frequently questioned. A “scientific policy” concern worth noting is the infrequency of course offerings: in response to this concern, the students have conducted a survey to collect data on what specific classes should be offered more regularly and what classes students have found most valuable.

The majority of explicit concerns that were expressed fall under “Resource Flows” and refer primarily to the intangible resources that result from a mentor/mentee relationship, rather than more traditional resources such as stipend. Even within their labs, graduate students have little control over resource flows and power dynamics (Sheltzer and Smith, 2014). Project distribution, hours, and authorship agreements are often informal, and there is little standardization of appropriate procedures and boundaries. As a result, students are subject to the supervisor’s decisions and changes in those decisions with little recourse (Mervis, 2016). The efforts we describe here that fall under the category of Resource Flows include a Bill of Rights which codifies some basic rights for students, as well as collecting regular data on resource distribution (e.g. demographics of award recipients) as called for by the 2007 National Academies report on women in STEM academia (Institute of Medicine, 2007). Formalizing a mechanism for enforcing student rights, clarifying the arbitration process, and adjusting resource distribution to meet program equity goals are all sensible and necessary explicit next steps.

Within the semi-explicit and implicit systems, nearly all of the concerns expressed in this exercise regarding relationships and power dynamics were “vertical” rather than “horizontal.” This indicates promotion of a positive peer-to-peer graduate student environment. But unfortunately, there is an extensive list of concerns surrounding the culture and infrastructure of mentorship. Our finding that graduate student concerns focus on dynamics between supervisor and mentee is unsurprising given that individual graduate experiences are often formed within the smaller context of the lab (Slay et al., 2019). Further, the academic system at predominantly white institutions does not inherently cultivate a culture of mentoring due to a focus on individual performance and independent productivity (Joseph, 2012; Meschitti and Lawton Smith, 2017). The nuanced role of an effective mentor includes scientific as well as psychosocial support (Lechuga, 2011). Effective mentorship has been empirically shown to be vital to graduate student success, particularly for under-represented groups (Riffle et al., 2013; Mack et al., 2013; San Miguel and Kim, 2015; for review see Makarem and Wang, 2020). The presence of a person the student considers a mentor who provides encouragement can counteract feelings of isolation experienced by women and under-represented minorities, as well as increase self-efficacy, job satisfaction, and work engagement (Tenenbaum et al., 2001; Handelsman et al., 2005; Meschitti and Lawton Smith, 2017). However, fear of appearing prejudiced has led to a “culture of silence” around the salience of race and ethnicity in mentor/mentee relationships in biology which prevents mentors from providing constructive psychosocial support for mentees (Byars-Winston et al., 2019). Our existing efforts to address these mentorship concerns focus broadly at the departmental level and are aimed at improving the student’s sense of support from the program, including a first year mentorship plan and personnel management training. Refocusing reform towards building accountability for quality of mentorship, rewarding effective mentors, and providing regular opportunities for mentorship skill building would target semi-explicit and implicit aspects of student concerns more directly and have a mutually beneficial impact on scientific outcomes (Lechuga, 2011; Varkey et al., 2012). For example, mentorship evaluation could be explicitly included in promotion and tenure applications and those who fall short could be asked to complete appropriate training.

The implicit concerns voiced by students in this exercise frequently and distinctly targeted women and minorities. Students were sometimes the subject of these incidents and frequently firsthand witnesses. Graduate programming to prevent or respond decisively to these interactions (jokes, comments, dismissal, or minimization) should be of the utmost importance. Our efforts to address implicit systems include a monthly discussion group for issues of equity and inclusion as well as trainings for students and faculty in bystander intervention and implicit bias. More regular and required training in restorative justice, anti-racism, personal resilience, decolonization, and anti-oppression would better match the regularity and pervasiveness students expressed (Bekki et al., 2013; Jackson et al., 2014).

However, implicit concerns, such as those regarding mental models such as biases, discrimination, and stereotype narratives, are highly intangible and societally pervasive. Paradoxically, the trainees who are most emphatically expressing these concerns are those most impaired by these mental models (Porter et al., 2018). Given this intangibility and inherent connection to power dynamics, change in explicit policies and practices is the primary available route for students to attempt to impact the implicit mental models of those with more seniority and power. We believe this creates an inherent mismatch within the systems change framework and hampers rectification on the implicit level. Therefore, faculty leadership in the ongoing work of altering mental models is vital. Policy change without progress in mental models, a core feature of a systems change framework, is not an effective tool for creating a more inclusive climate.

The circulation and presentation of the 2018 departmental climate survey was critical in changing mental models regarding the pervasiveness and seriousness of graduate student concerns (see supplemental material). In the survey, students and staff were asked to reflect on their experiences and evaluate how included they felt in various aspects related to their identity. Similar surveys have been implemented in other departments (Princeton Graduate Women in STEM Leadership Council, 2018). Cross-hierarchical communication has the ability to “change the narrative” and is a crucial step in long-lasting systemic change.

## Conclusions

The academic system is touted as a meritocracy, but in reality, it still embraces norms and policies that are inequitable and paternalistic, leading to lower productivity and job satisfaction (Settles et al., 2006). When diverse perspectives are lost from the academic system, the quality, scholarship, and innovation of the institution is diminished (Østergaard et al., 2011; Díaz-García et al., 2013; Adams, 2013; Freeman and Huang, 2014; AlShebli et al., 2018). The loss of competitive colleagues and the dampening of academic aspirations occurs, not for scientific reasons, but due to a lack of salient support for the types, and disproportionate number, of barriers experienced by women and under-represented minorities (Handelsman et al., 2005; Riffle et al., 2013; Ong et al., 2011; Mack et al., 2013). Pushing back on these inequities by supporting graduate student trainees is vital to generating a more just scientific enterprise. Improving the experiences of graduate students, particularly marginalized students, is vital to our collective publication fecundity, effective teaching, competitive recruitment, prolific grant applications, and high quality research (Adams, 2013; Freeman and Huang, 2014; AlShebli et al., 2018).

In many cases there is an initial energy by graduate programs to improve inclusion, often led by individual students perceptive to these systemic challenges (Porter et al., 2018). But addressing complex issues that ultimately stem from deep societal iniquities and power structures cannot be solved in a single hour-long bias training event (Jackson et al., 2014). Efforts must be multimodal, present at all levels of the systems change framework, and consistent to be effective.

Graduate students lack power to generate systemic change not only because of limited financial resources and institutional power, but also because of the loss in institutional knowledge that occurs as students graduate, and no mechanism exists for systematically passing knowledge to new students. As discussed, this is part of why a systems change framework makes the practice of critical evaluation more effective. However, concerns and efforts initiated by graduate students must be respected and maintained by more permanent members of a program such as faculty and staff. Additionally, department wide conversations, events, and trainings should be held regularly to allow ample opportunity for communications and interactions between levels of the hierarchy. As we describe here, a critical first step in this process is to evaluate your own programs’ diversity, inclusivity, and climate efforts to determine how well they reflect graduate student concerns is a critical step in implementing effective change.

In assessing our efforts, we acknowledge our limitations of our mental models, experiences, and, at times, reliance on anecdotal data. We hope that our practice spurs more comprehensive exercises of this nature and research into effective systems change within academia. Future exercises should include regular re-evaluation across trainee cohorts after policy implementations, to gauge how the program needs and improvements grow and change over time. Comparisons between similar programs could stimulate compelling and useful cross-institutional discussion. Future work should focus increasingly on the intersectionality of identities, challenges, and experiences within the academic system.

We hope that the pedagogical exercise constructed here can help programs critically assess their efforts to improve climate and culture, and we hope this in turn positively impacts diversity and inclusivity in both direct and indirect ways. Program efforts that thoughtfully and actively enrich trainee experiences in graduate school promote a healthier and happier scientific community.

## Supporting information

Electronic Supplementary Materials

## Acknowledgements

The authors would like to thank the students who participated the climate survey and participated in discussions with the authors. We would like to thank Marian Schmidt and Brian Barnes for manuscript editing, and two anonymous reviewers for comments and support.

## Data Accessibility Statement

Primary data related to Figure 1 can be obtained by contacting.

## Competing Interests Statement

The authors declare no competing interests.

## Author Contributions

KW conceived of the report. JY collected quantitative data. KW and JY wrote the manuscript. All authors give final approval for publication.

## Notes

### Competing Interest Statement

The authors have declared no competing interest.

