## Supplementary material for "A Systems Change Framework for Evaluating Academic Equity and Inclusion in an Ecology & Evolution Graduate Program": Electronic Supplementary Materials

Supplementary Materials - EEB Grad Student Efforts + Systems Change
Kelly Wallace & Julia York

Table of Contents

1. Data Request Form
2. Department of Integrative Biology Climate Survey
3. EEB Program Graduate Student Bill of Rights
4. EEB First Year Mentoring Plan
5. Information on gender neutral bathrooms and quiet/lactation rooms
6. EEB Course Offering Survey

**Supplemental Material 1**: Data Request Form

---

We used the following form to request data on the department and graduate program from the College-level Dean's office, the departmental administration, and the program coordinator. Not all the data requested is collected and not all the data was provided. However, this can serve as a starting point for assessing the current representation, resource distribution, recruitment, and career success of program/department members and can be adapted for use in other contexts.

**Equity and Inclusivity**

Department of Integrative Biology

Goal: we plan to gather statistics on representation and identity among employees and students in the Department of Integrative Biology at UT Austin. We are motivated to assess the current state of the Department in terms of representation across the hierarchy, distribution of resources, recruitment and retention, and equality in career success whether one remains at UT or elsewhere. To that end, we aim to gather identity information for the following groups.

**Identity information for each group:** gender, race, disability, years since PhD, immigration status

**Representation:**

All faculty

Full professors

Associate professors

Assistant professors

Adjunct professors

Graduate students

Postdocs

Staff

Faculty service committee members

Faculty administrators

Invited speakers

Speaker hosts

**Distribution of resources:**

Faculty salary

Postdoc salary

Staff salary

Graduate students holding fellowships

Number of TA semesters for graduate students

**Retention:**

Time of employment for faculty

Time of employment for staff

Time to graduate for students

Students receiving PhDs

Students received Masters

Students dropping out

Faculty hired elsewhere

Faculty quitting

Staff quitting

**Recruitment:**

New faculty hires over the last 5 years

Starting salary

Start-up package

Graduate students offered admission

Graduate students accepting admission

Graduate students applying

**Equality in career success:**

Faculty receiving tenure

Average time to tenure

Average number of promotions

Average time to promotion

Graduates employed upon graduation

Graduates employed in academia

Graduates employed in industry

Postdocs receiving faculty jobs

Average postdoc positions

Average weekly time spent on service

Average weekly time spent teaching

Average weekly time spent on research

Numbers of papers published

**Supplemental Material 2**: Department of Integrative Biology Climate Survey

---

### IB Cultural Climate Survey

If you are at all affiliated with or have experience with the Department of Integrative Biology at UT Austin, please complete the form. We have added questions to categorize responses, but if you prefer not to give any identifying information please choose "prefer not to say." Any identifying information will be confidential.

NOTE: ALL RESPONSES ARE ANONYMOUS. PLEASE DO NOT INCLUDE IDENTIFYING INFORMATION IN YOUR RESPONSE. PLEASE USE GENERAL IDENTIFIERS SUCH AS "MY ADVISOR", "A PROFESSOR", OR "A GRAD STUDENT." If you have an incident that you would like to report in a more detailed manner, please use this form: https://goo.gl/forms/jKz6C7uKuumGUA9j1

IMPORTANT: We are not collecting this information for reporting purposes. IF YOU NEED TO REPORT AN INCIDENT, PLEASE CONTACT THE TITLE IX OFFICE (http://titleix.utexas.edu/), THE CAMPUS CLIMATE RESPONSE TEAM (http://diversity.utexas.edu/ccrt/), OR THE OFFICE OF INCLUSION AND EQUITY (http://equity.utexas.edu/). We are collecting this information to get an empirical evaluation of experiences in Integrative Biology.

Any use of this information will be anonymous.

Are you affiliated with the Integrative Biology (IB) department at UT Austin?

- Yes
- No
- Other:

What is your role in the department?

- Graduate student
- Postdoc
- Staff
- Faculty
- Prefer not to say
- Other:

With what gender do you identify?

Female

Male

Nonbinary

Prefer not to say

Other:

Do you feel welcome in the department and at departmental events?

Not at all

1

2

3

4

5

Yes, completely

Are you satisfied with the overall culture of the department?

Not at all

1

2

3

4

5

Yes, completely

Do you feel valued in the department?

Not at all

1

2

3

4

5

Yes, completely

Please provide any further information on your answers in this section and indicate if you are comfortable having it anonymously shared with faculty.

**Supplemental Material 3**: EEB Program Graduate Student Bill of RIghts (October 2019)

---

*Background Info: This document should clarify what graduate students are entitled to and should expect from their advisor(s), the EEB program, and the IB department. We have limited this bill of rights to items that are within advisor(s) and/or the IB department's control. For example, given that wages, disability accommodations, and maternity/paternity leave, are rights that are determined by the University, they are beyond the scope of this bill. Furthermore, determining how item 10) will be carried out and whom students should contact to report a violation is also beyond the scope of graduate student input and will be left up to the faculty to determine.*

---

(1) The right to be treated without bias with respect to (but not limited to) gender, race, age, sexual orientation, gender expression, disability, religious or political affiliations, family status, country of origin, mental/physical health, and citizenship.

(2) The right to a clear understanding of the responsibilities of a graduate student to their advisor(s).

(3) The right to mentoring from their advisor(s) that supports and advances the student's academic and professional goals.

(4) The right to clear, concrete minimum requirements for graduation (masters or PhD). NOTE: A section should be added to the graduate handbook that includes these minimum requirements

(5) The right to explore or pursue a career outside academia without penalty. Specifically, plans to pursue a non-academic career after graduation will not be grounds for denying financial or supervisory support, either by the advisor or the program.

(6) The right to meet the standard hourly contract (both RA and TA positions) and to stay within this contract.

(7) The right to take courses that promote a student's career goals and to access professional training courses and seminars as needed.

(8) The right to fair/accurate representation of a student's work and contributions.

(9) The right to have a student representative to advocate and represent EEB graduate student interests at faculty meetings.

(10) The right to a non-biased arbitration process if and when seeking to resolve a violation of these rights. Official academic grievance procedures designed and administered by the University and the Graduate School are beyond the scope of this document. Supplementary, less formal complaint procedures should be clearly specified by the IB department and by the EEB graduate program, and clearly communicated to to graduate students at the time of entry.

(11) The right to be informed of these rights upon enrollment, and to be free of reprisals for exercising these rights.

**Supplemental Material 4**: EEB Program First Year Mentoring Plan

---

*Overview:*
*The EEB Graduate Program Mentoring Plan is a form designed to provide incoming EEB Graduate students (mentees) and their PI's (mentors) an opportunity to outline goals, expectations, and strategies for a productive mentor-mentee relationship.*

*The Mentoring Plan is a form split into two main sections: an initial meeting and a follow-up meeting. The mentor and mentee are encouraged to think about the questions in the form before the initial meeting and seek guidance from peers. The mentor and mentee should then meet in person during the mentee's first semester to co-create and print out a document that answers the questions provided. The follow-up meeting should take place in the student's second semester.*

*This plan is flexible, and it is encouraged that even beyond the two required meetings the mentor and mentee can continue to refer to the document, update it, and use it as a framework to engage in productive communication about mentorship.*

*This mentoring plan utilizes components of the Stanford Biosciences Individual Mentoring Plan (Year 1) https://biosciences.stanford.edu/wp-content/uploads/2018/01/IDP_Year_1.pdf*

*Part 1: Initial Meeting*

**Overview (mentor):** What is important to you, mentor, in a mentoring relationship?

**Overview (mentee):** What is important to you, mentee, in this mentoring relationship?

**Communication (frequency):** How often will the mentor and mentee meet?

**Communication (preferences):** What is the best way for the mentor and mentee to communicate regularly?

**Expectations (academic):** What are the academic interests of the mentee? These can be keywords, questions, project ideas, etc.

**Expectations (funding):** What is/are the mentee's expected funding source(s) for this current year (and in the future if known)? What resources are available for the mentee to pursue funding opportunities?

**Expectations (responsibilities):** What are the responsibilities of the mentee (including e.g. training, lab operations, time commitments)

**Goals (requirements):** What EEB program requirement goals does the mentee plan to accomplish by the end of this year? Are there any strategies the mentor or mentee can use to accomplish these goals?

**Goals (research):** What research goals does the mentee plan to accomplish by the end of this year? Are there any strategies the mentor or mentee can use to accomplish these goals?

**Skills (research):** What skills (pick 2, listed below) does the mentee identify as important development targets for the coming year? How will the mentor and mentee develop these areas?

**Skills (communication):** What skills (pick 2, listed below) does the mentee identify as important development targets for the coming year? How will the mentor and mentee develop these areas?

**Research Skills:**
Broad-based knowledge of science
Critical reading of scientific literature
Experimental design
Statistical analysis and interpretation of data
Creativity and innovative thinking
Understanding of submission/peer review process
Identifying and seeking advice
Time management

**Communication Skills:**
Writing for a research proposal or publication
Writing with appropriate grammar and structure
Speaking to a specific audience
Communicating one-on-one
English fluency
Working with constructive criticism

**Challenges:** What is the mentee's main concern regarding their transition into graduate school and setting themselves up for success in their PhD? How can the mentor help navigate those concerns?

**Challenges:** How do the mentor and mentee plan to address any issues (e.g.miscommunication, conflict) that arise?

**Other:** What other information and/or resources should the mentor be aware of to help foster a productive mentor-mentee relationships?

**Other:** What other information and/or resources should the mentee be aware of to help foster a productive mentor-mentee relationships?

**Follow- Up:** Please schedule a date for a follow-up meeting during the mentee's second semester.

*Part 2: Follow-Up Meeting*

**Goals:** What has the mentee accomplished in their first semester?

**Communication:** How has the communication between the mentor and mentee been this past semester/ How could it improve?

**Expectations:** Have the mentee's research interests changed since the initial meeting?

**Skills (accomplishments):** How has the mentee tackled improving any of the skills listed below?

**Skills (development):** For the specific skills highlighted during the initial meeting as targets for development, has any progress been made to develop those skills? If so, how? If not, how can they be made a priority in the future?

**Follow- Up:** How can the mentor and mentee continue to improve their relationship? Would the mentor or mentee like an additional follow-up meeting?

**Research Skills:**
Broad-based knowledge of science
Critical reading of scientific literature
Experimental design
Statistical analysis and interpretation of data
Creativity and innovative thinking
Understanding of submission/peer review process
Identifying and seeking advice
Time management

**Communication Skills:**
Writing for a research proposal or publication
Writing with appropriate grammar and structure
Speaking to a specific audience
Communicating one-on-one
English fluency
Working with constructive criticism

**Supplemental Material 5**: Information on gender neutral bathrooms and quiet/lactation rooms

---

Gender neutral bathroom options should be provided for the use of employees and students who do not identify with a binary gender or who feel unsafe in selecting a gendered bathroom. Often this is simply a matter of changing signage. The College of Natural Sciences at the University of Texas at Austin's policy is now that all single stall bathrooms must be signed as gender neutral or gender inclusive. More information can be found at: https://diversity.utexas.edu/genderandsexuality/gender-inclusive-restrooms/

Quiet and Lactation rooms provide clean, quiet, and private space for nursing mothers. However, they are also for the use of those who require space to rest, such as those with non-contagious medical conditions. Ideally, this space has a sink, multiple electrical outlets, refrigerator, comfortable chair, paper towels, breast pump, hand sanitizer, a lamp, a clock, a mirror, and soap. For more information on Quiet and Lactation Rooms visit:

**Supplemental Material 6**: EEB Course Offering Survey

---

*This survey was administered March 2019.*

1. Did you take coursework or formal training in Ecology before joining the program?
2. Did you take coursework or formal training in Evolution before joining the program?
3. Did you take coursework or formal training in Behavior before joining the program?
4. Did you take coursework or formal training in Statistics before joining the program?
5. Was there any other relevant coursework you took prior to joining the program that you feel is relevant to share? (please leave blank if not)
6. How satisfied are you with the FREQUENCY of graduate course offerings?
7. How satisfied are you with the NUMBER of courses offered?
8. How satisfied are you with the coursework-related graduation REQUIREMENTS?
9. How satisfied are you with the graduate coursework OVERALL?
10. How would you describe the purpose of the first year core course that you took?
11. What do you think the purpose of the first year course SHOULD be?
12. What TOPICS/CONTENT do you think should be included in the first year course?
13. What SKILLS do you think should be included in a first year course?
14. How satisfied are you with the CONTENT of your first year EEB course?
15. How satisfied are you with how the first year course was taught?
16. What are some courses that you have really enjoyed in your graduate coursework?
17. What are some courses that you feel could have been more useful than they were? Why?
18. What topics/content/skills do you wish could be offered as a course?
